# Infectivity of dengue virus serotypes 1 and 2 is correlated to E protein intrinsic dynamics but not to envelope conformations

**DOI:** 10.1101/303560

**Authors:** Kamal Kant Sharma, Xin-Xiang Lim, Sarala Neomi Tantirimudalige, Anjali Gupta, Jan K. Marzinek, Daniel Holdbrook, Xin Ying Elisa Lim, Peter J. Bond, Ganesh S. Anand, Thorsten Wohland

**Author notes:** To whom correspondence should be addressed: Thorsten Wohland, Centre of bioimaging Sciences, Department of Biological Sciences, National University of Singapore, 14 Science Drive 4, Singapore 117543, (65) 6516 1248.

## Abstract

Dengue is a mosquito-borne virus with dire health and economic impact. Dengue is responsible for an estimated ~390 million infections per year, with Dengue 2 (DENV2) being the most virulent strain among the four serotypes. Interestingly, it is also for strains of this serotype that temperature-dependent large scale morphological changes, termed as “breathing”, have been observed. Although, the structure of these morphologies has been solved to 3.5 Å resolution, the dynamics of the viral envelope are unknown. Here, we combine fluorescence and mass spectrometry and molecular dynamics simulations to provide insights into DENV2 structural dynamics in comparison to DENV1. We observe hitherto unseen conformational changes and structural dynamics of the DENV2 envelope that are influenced by both temperature and divalent cations. Our results show that for DENV2 and DENV1 the intrinsic dynamics but not the specific morphologies are correlated to viral infectivity.

Graphical Abstract/ cover Figure

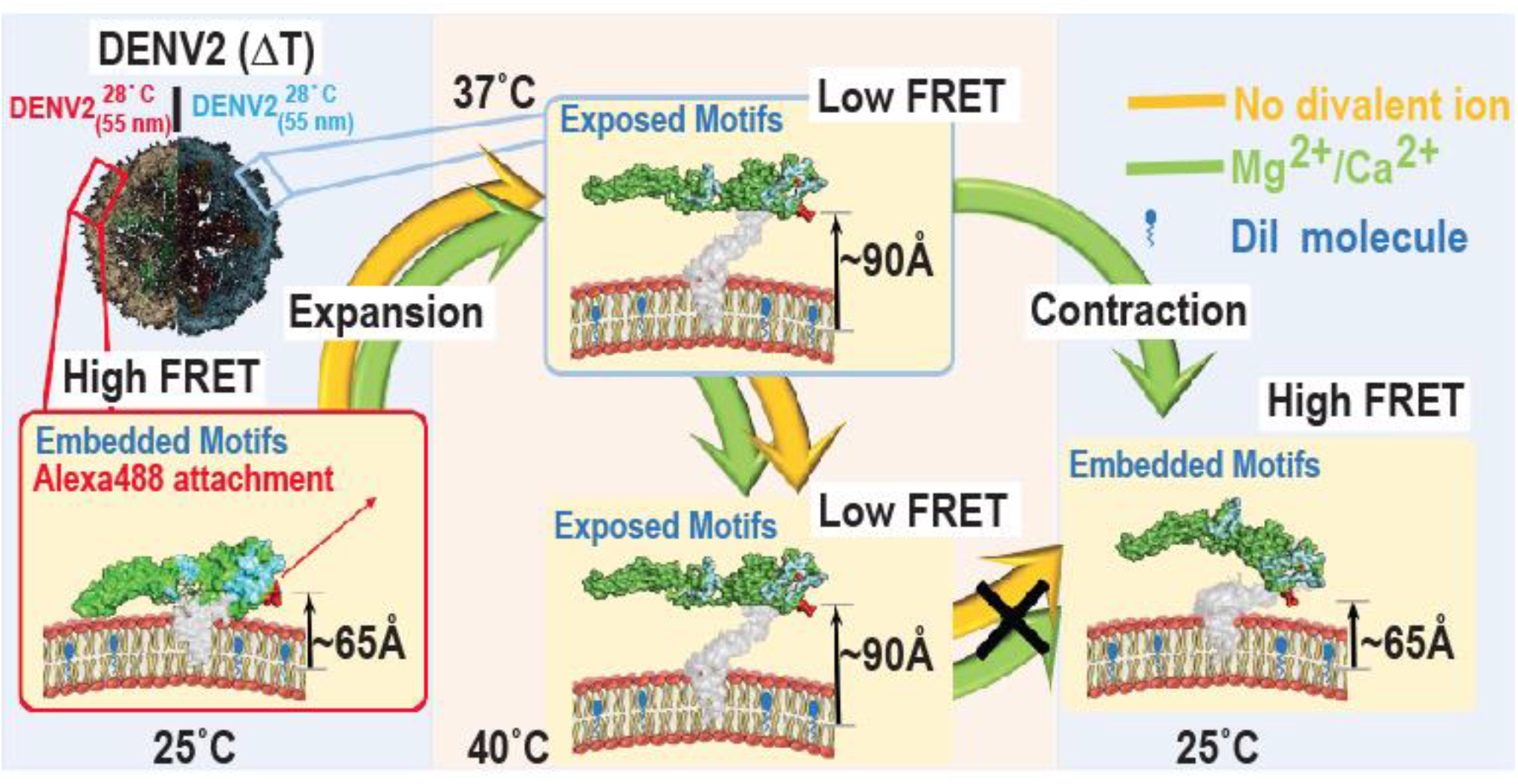

## Introduction

Dengue Virus (DENV) is responsible for a wide range of clinical manifestations ranging from acute febrile illness to life-threatening dengue haemorrhagic fever (DHF) and dengue shock syndrome (DSS) (1–4) with an estimated ~390 million infections annually (5). DENV has four characterized serotypes (DENV1-4) and infection by one serotype does not confer immunity against heterologous serotypes (6). Instead, it may cause a more severe and life-threatening form of dengue infection through antibody-dependent enhancement (7–9). In spite of such a disease burden, there are no alternative therapeutics (9–12) with the exception of a few low efficacy (~30% overall) vaccines (9, 12) in the market. Evidence suggests that flaviviruses explore an ensemble of morphologies at equilibrium, referred to as viral structural dynamics or “breathing” (13–16) and may be an essential aspect of the life cycle of many viruses (13, 17). These distinct morphologies arise due to large and small scale conformational changes in the organization of surface glycoproteins which results from the virus experiencing perturbations in temperature, pH or host-protein interactions (18). DENV(1–4) display a variable degree of surface glycoprotein (E protein) organization at temperatures above 34°C, as in a human body (37°C), than they do at lower temperatures, such as those found in mosquitoes (28°C) (19–23). Temperature dependent transitions in E protein organization are most prominent in DENV2 which acquires a “bumpy” form at temperatures above 35°C in contrast to the “smooth” form at 28°C (15). It has been suggested that such large amplitude reversible morphological changes in the DENV2 envelope might be the basis of “breathing”, which could be blocked by the insertion of an antibody (15). However, the formation of the DENV2 bumpy form has been observed to be irreversible, at least for two different strains, DENV2 16681 and DENV2 New Guinea Strain (NGC) (20, 23). Furthermore, plaque assays indicated that the smooth and bumpy morphologies are not correlated with infectivity (20). This raises at least two questions regarding DENV conformations, dynamics and infectivity. First, are the morphological changes really irreversible? Structural experiments have been conducted in buffers that lack divalent ions contrary to the plaque assays, which are conducted in cell culture media in the presence of Ca^2^+ and Mg^2^+. As the structure and function of several viruses depends on divalent ions (24–27), is it possible that DENV conformations also depend on divalent ions? Second, is it possible that there exist viral protein dynamics at faster time and smaller length scales, as suggested by Zhang et al. (23) that are related to infectivity? For the sake of clarity, in this article we will distinguish between DENV morphologies, which are induced by temperature changes and are accessible by electron microscopic structural investigations (to date called breathing), and DENV intrinsic dynamics, which are fast protein dynamics at smaller amplitudes and which are not related to any persistent structures (Figure 1).

**Figure 1:**
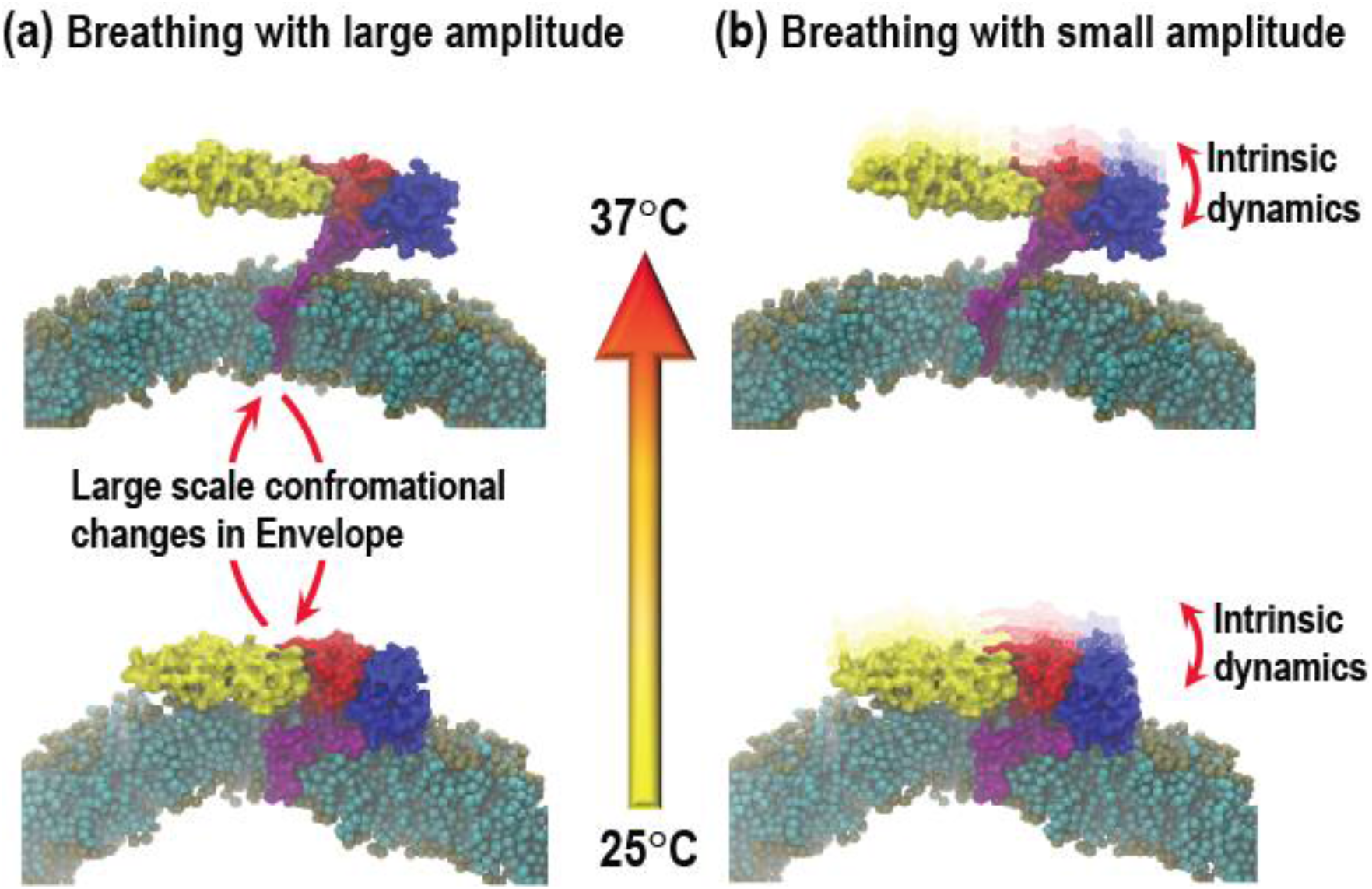
Schematic representation of DENV2 “breathing” with (a) large and (b) small amplitude. **(a)** DENV2 undergoes temperature dependent large scale conformational changes in the organization of surface E proteins between 25°C and 37°C. In smooth DENV2, the E protein lies close to virus bilayer. While in bumpy DENV2, the E protein undergo outward motion and a 15 Å gap between lipid bilayer and E proteins layer is formed that in turn results in the expansion of virus from 50 nm diameter (at 28°C) to the diameter of 55 nm (at 37°C) (19, 20). However, unlike depicted in scheme, the lowering of temperature does not induce reversal of large scale conformational changes in E protein and thus defying the concept of DENV2 “breathing”. **(b)** DENV2 breathing also includes the E protein structural dynamics that takes place in both smooth and bumpy form at smaller time and length scales (23), termed as intrinsic dynamics.

We have earlier followed the large scale outward conformational motion of envelope proteins (expansion) in DENV2 and DENV1 between 25°C and 40°C by time-resolved Förster Resonance Energy Transfer (trFRET), using dual-labeled virus particles that carried Alexa488-TFP as FRET donor on the E protein and DiI as FRET acceptor within the lipid bilayer (22) (Figure 2a). In trFRET, the energy transfer from donor to acceptor, which is strongly distance dependent, influences the donor fluorescence lifetime. The Förster radius, *R_0_*, of the pair is 59 Å, which is on the order of the distance between E protein and bilayer (20). The large scale morphological changes of DENV alters the distances between the E protein and the viral lipid membrane (28) and in turn, influences the donor fluorescence lifetime. However, we did not investigate the temperature dependent reversibility (contraction) of those morphological changes in dependence of divalent ions, nor did we investigate fast low amplitude dynamics. Here, we validated that the formation of the DENV2 bumpy form is irreversible in absence of divalent cations. For this, we used DENV2 (NGC) and DENV1 (PVP 159) (DENV2 and DENV1 in the rest of the article refer specifically to NGC and PVP159 strains, respectively), produced and purified from C6/36 *aedes albopictus* mosquito cells at 28°C. We further investigated, if these temperature dependent large scale morphological changes are reversible in presence of divalent cations.

**Figure 2:**
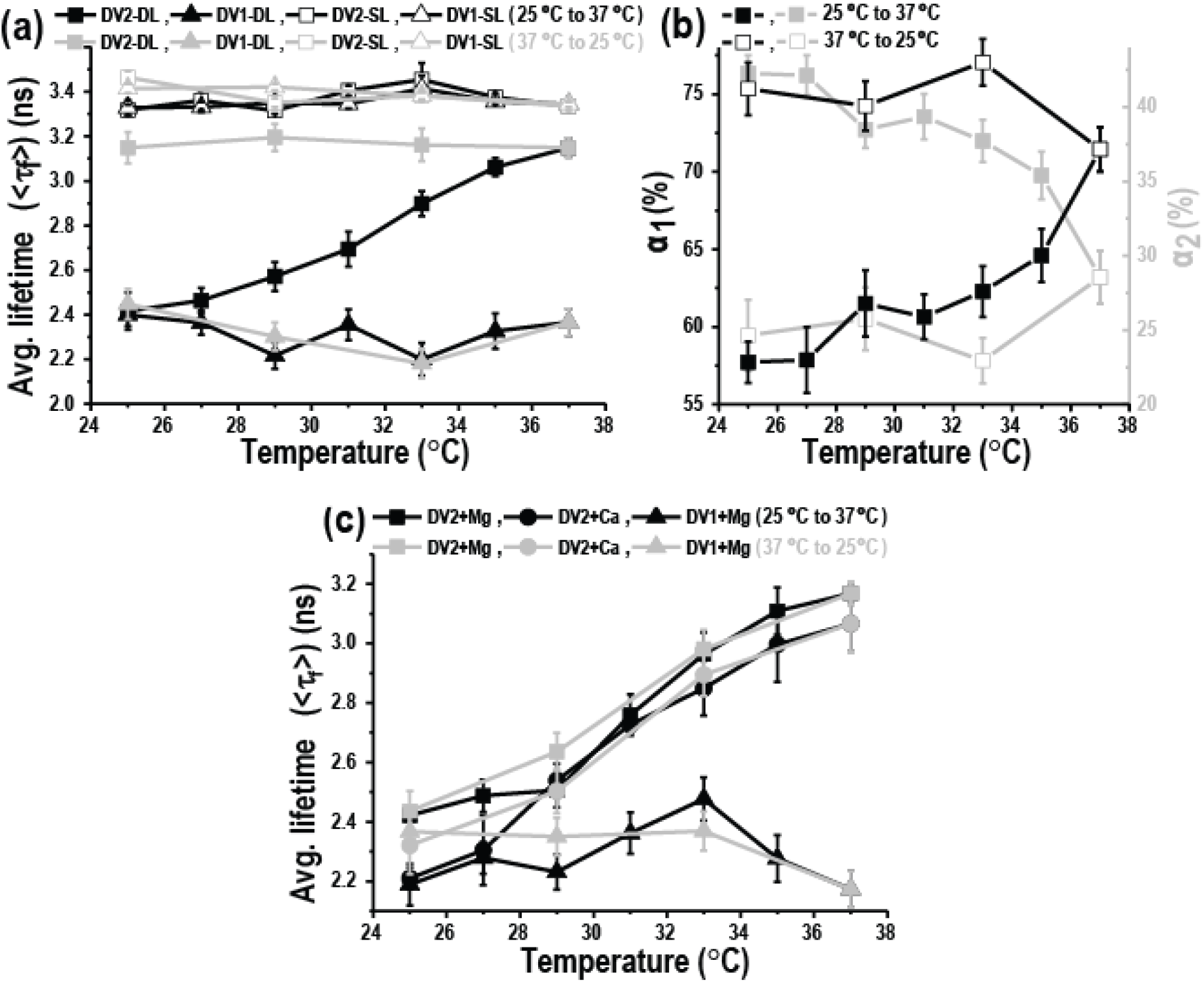
Structural transitions of DENV2 and DENV1 between 25°C and 37°C in (b) absence and (c) presence of Mg^2+^/Ca^2+^-ions. The lifetime traces for single and dual labeled viruses were fitted to mono-and bi-exponential decay model, respectively. **(a)** The average fluorescence lifetimes (<τ_f_>) of AF488-TFP for donor only labeled, <τ_f, SL_> (open triangles/squares) and dual labeled, <τ_f, DL_>, (filled triangles/squares) for 5 × 10^7^ pfu/ml DENV2 (squares) and 5 × 10^7^ pfu/ml DENV1 (triangles). The data was collected during temperature increase from 25°C to 37°C (black triangles/squares) followed by decrease from 37°C to 25°C (grey triangles/squares). **(b)** Lifetime amplitudes corresponding to populations with low-(black open/filled squares) and high-FRET (grey open/filled squares). The data was collected during temperature increase from 25°C to 37°C (filled squares) followed by decrease from 37°C to 25°C (open squares). By fitting dual labeled viruses to a bi-exponential decay model, two lifetimes were calculated. The value of ~3.7 ns corresponds to unquenched or expanded DENV2, whereas the second lifetime of ~1.2 ns represents DENV2 particles that were unexpanded or undergo contraction. **(c)** The average fluorescence lifetimes (<τ_f_>) DENV2 in presence of 1 mM Mg^2^+-ions (squares) or 2 mM Ca^2^+-ions (circles) and for DENV1 in presence of 1 mM Mg^2^+ions (triangles). The data was collected during temperature increase from 25°C to 37°C (black circles/triangles/squares) followed by decrease from 37°C to 25°C (grey circles/triangles/squares). Error bars represents standard deviations of six different experimental replicates in both DENV2 and DENV1 viruses.

In addition to large scale morphological changes, we also followed intrinsic dynamics of the single DENV envelope protein by Förster Resonance Energy Transfer fluctuations spectroscopy (FRET-FCS), both in presence and absence of divalent cations. FRET-FCS uses the same labeled virus but with much higher time resolution to investigate the intrinsic dynamics on the millisecond time scale. Thus, the labeling with donor and acceptor fulfills two tasks: (i) monitoring of viral morphological changes and (ii) measuring the protein dynamics and flexibility.

We supported the fluorescence results with amide hydrogen/deuterium exchange mass spectrometry (HDXMS), molecular dynamics (MD) simulations, and infectivity measurements via plaque-forming assays. HDXMS captures thermodynamic transitions of proteins by measuring the differences in mass when backbone amide hydrogens exchange with solvent deuterium (29, 30). The extent of HDX is dependent on hydrogen bonding and solvent accessibility of the protein (31, 32), without the limitation of size/quaternary assembly (33), at the second and slower time scales (31). In addition, MD simulations (34–36) and coarse-grained (CG) methodologies enable access to dynamics (37–39) while plaque assay measures the infectivity of a virus (40, 41). Taken together our results show that, at least for DENV2 and DENV1, viral intrinsic dynamics but not the specific morphologies correlate with DENV infectivity.

## Results and Discussion

### FRET-pair labeled DENV1 and DENV2 show temperature dependent lifetimes

DENV2 E protein, constituting the viral envelope, has four domains (E-DI (residues 1-52; 132-193; 280-296), E-DII (residues 53-131; 194-279), E-DIII (residues 297-394) and the stem domain (residues 395-486)) that connect to a transmembrane region (42, 43). The pseudo-atomic resolution map of the virus capsid (44) showed that the E protein forms 90 dimers on the smooth surface of the mature virus (20). Cyro-electron microscopy (20, 23) (cryo EM) and HDXMS (22) studies revealed that these E protein dimers undergo an outward motion during a temperature change from 28ºC and 37ºC, leading to a morphological expansion from a “smooth” to a “bumpy” form of the virus. In buffer containing 150 mM sodium ion solution, this temperature dependent morphological change was irreversible even when the temperature was subsequently lowered to 28ºC. As stated earlier, by investigating the outward motion of E proteins from the virus bilayer for DENV2 and DENV1 using trFRET(22), we confirmed the temperature dependent morphological changes, due to virus expasion, seen by EM and HDXMS. However, the regions in E protein that are responsible for the donor lifetime changes in trFRET, were missing.

Since, Alexa488-TFP attack primary amines, the labeling site on the E protein will depend on the accessibility of the lysine and its N-terminus. To identify the labeled regions, we tagged the viruses with N-hydroxysulfosuccinimide ester of biotin, which labels free amino groups, similar to the fluorescence tag, Alexa488-TFP. The biotinylated peptides were captured using high affinity biotin-streptavidin interactions. The position of biotinylated residues was determined by peptide mapping using LC-MS/MS. The majority of biotins were found coupled to the lysine residue (K344) in the region spanning peptides 335_I_-345_V_ (Figure S1a) of the DENV2 E-DIII. K344 in the motif 335_I_-345_V_ is proximal (<10 Å) to the DENV2 E protein peptide residues 307_V_–338_K_ (denoted by orange color in Figure S1a) that undergo outward conformational changes between 28°C and 37°C (22). This suggests that the trFRET monitored temperature dependent large scale DENV2 morphological changes represents the large scale conformational changes in DENV2 E-DIII.

Similarly, for DENV1, the majority of biotins were found coupled to the lysine residue (K110 and K135) in the region spanning peptides 95_T_–110_K_ and 129_I_–136_Y_ (denoted by red color in Figure S1b) of DENV1 E-DII. The motifs 95_T_–110_K_ and 129_I_–136_Y_ are in proximity to the DENV1 E protein peptide residues 238t-260h (DENV1 E-DII) and 152_N_–173_Q_ (DENV1 E-DI), respectively (denoted by orange color in Figure S1b) that undergo conformational changes between 37°C and 40°C (22).

As control measurements, we measured donor-only labeled virus at 25ºC which could be fitted with a single lifetime of 3.47 ± 0.07 ns for both DENV1 and DENV2 (Figure 2a). The donor and acceptor labeled viruses exhibited two lifetimes at the same temperature. The decay of DENV2 had an average lifetime of 2.42 ± 0.05 ns (Figure 2b) and was best fitted with two discrete (long and short) lifetime components of 3.72 ± 0.1 ns (<τ_1_>) and 1.18 ± 0.1 ns (<τ_2_>) (Figure S2b) having populations of 57 ± 2 % (α1) and 43 ± 2% (α2) (Figure 2a), respectively. The donor fluorescence intensity decays of DENV1 had an average lifetime of 2.37 ± 0.1 ns (Figure 2b) and were also best fitted with two discrete lifetimes with values similar to DENV2 (3.7 ± 0.1 ns (<τ_1_>) and 1.14 ± 0.1 ns (<τ_2_>)), with α1 and α2 measured as 46 ± 2 % and 54 ± 2 %, respectively (Figure S2e), indicating viral structural heterogeneity. The fact that the lifetimes for dual-labelled viruses are significantly shorter than the average lifetimes for corresponding single-labeled viruses indicates the existence of FRET at a population averaged distance of ~65 Å (Figure S1c) between virus lipid bilayer and its E protein scaffold. Interestingly, these discrete lifetimes (<τ_1_> and <τ_2_>) remained constant over the full temperature range from 25ºC and 40ºC, and only the population (α_1_ and α_2_) of molecules corresponding to these lifetimes showed changes. Since our data contains only two populations and an increase in the value of α_1_ will lead to a decrease in the value of α_2_ by the same amount, we report in the rest of the article the population corresponding to only the long-lifetime (α1) and the average lifetime. Data for individual lifetimes are provided in the supplement.

### DENV2, but not DENV1, shows Mg^2^+/Ca^2^dependent morphological changes between 25ºC and 37°C

In the past DENV morphological expansion had only been investigated in aqueous solutions containing only monovalent ions (20, 22, 23, 45). However, the molecular dynamics that led to this expansion were not addressed raising several questions. Are all viral particles expanding equally? Are the morphological changes reversible, i.e. are the viral particles in thermal equilibrium?

First, as control, we confirmed the morphological expansion of the DENV 1 and DENV2 during the transition from 25°C to 37°C. Then, we investigated reversibility of the morphological changes when the temperature was lowered back to 25°C, similar to the transition between humans and mosquitos.

An increase in temperature from 25°C to 37°C in steps of 2°C led to an increase of the average fluorescence lifetime, <*τ*_f_>, (2.42 ± 0.05 ns → 3.18 ± 0.05 ns) and the long-lifetime population, α1, (57 ± 2 % → 72 ± 3 %) for DENV2 (Figure 2a & 2b). The increase in <*τ*_f_> indicated that the average distance between the labeled E-DIII (307_V_–338_K_) and viral membrane bilayer increased from ~65 Å to ~90 Å (Figure S1c). While the difference in the values of α1 at 37°C and 25°C indicated only ~15% from the total labeled DENV2 viruses underwent morphological transitions. This heterogeneity in DENV2 structures is consistent with literature, where only ~22% of DENV2 particles show homogeneity in virus morphology (23). A similar structural heterogeneity is reported for other flaviviruses (13). However, in addition to the E-DIII peptide region 307_V_–338_K_, HDXMS has mapped other DENV2 E protein motifs that undergo conformational changes during expansion from 28°C to 37°C(22) viz., peptides spanning residues from E-DII (78_G_–107_F_ and 238_T_–278_F_), E-DI (21_V_–30_C_), the second and third stem helices (431_I_-448_S_) and transmembrane helices (465_I_–486_L_) (22). With stepwise decrease in temperature from 37°C to 25°C, a marginal or no change in the values of <*τ*_f_> and α_1_ indicates no changes in the average distance between the labeled E-DIII (307_V_–338_K_) motif and the viral membrane (Figure 2b & Figure S1c). This result is in line with literature (20, 22, 23, 45) and confirms the irreversibility of temperature dependent DENV2 morphological changes in absence of divalent cations.

The introduction of divalent cations (Mg^2+^ and Ca^2+^) does not significantly alter the population that undergoes structural transitions (Figure S2c and S2d). In addition to this, the similar values obtained for <*τ*_f_> either at 25°C or 37°C (Figure 2c), suggest that divalent cations do not interfere with the viral heterogeneity. However, with stepwise reversal in temperature from 37°C to 25°C, a decrease in values of the <*τ*_f_> (~3.2 ns → ~2.3 ns) (Figure 2c) and α1 (~72 % → ~58 %) (Figure S2c and S2d) were observed for DENV2. This reversal in <τ_f_> values (~2.4 ns → ~3.2 ns → ~2.3 ns) suggests that the irreversibility of temperature dependent DENV2 morphological changes is partly due to the absence of divalent cations.

Surprisingly, only 1 among the 6 E protein motifs (22), namely the E-DIII region spanning peptides 307_V_–338_K_, showed temperature dependent reversal in deuterium exchange when reverting to 28°C as compared to deuterium exchange at 37°C (Figure 3a). However, this conformational reversal of peptide 307_V_–338_K_ was observed to be independent of the Mg^2^+ions.

Furthermore, unlike peptide 307_V_–338_K_, the E-DIII region spanning peptides 378_I_-391_F_ showed Mg^2^+ion dependent negative deuterium exchange when reverting to 28°C (Figure 3a). The results showed that only two E-DIII motifs undergo temperature dependent reversible large scale conformational changes and thus indicates a minimal change in the DENV2 bumpy morphology when reverting to 28°C (Figure S3). This is in contrast to picornavirus like “breathing” where viruses undergo temperature dependent large scale reversible conformational changes (17, 46–49).

Thus, in addition to establishing E-DIII flexibility during both outward and inward conformational changes, these results also prove that DENV2 undergoes only partial “breathing” and that only in presence of divalent cations. However, such partial DENV2 morphological changes should not be neglected as E-DIII is involved in host-receptor binding (42, 43, 50, 51). And it is possible that such divalent cation dependent large scale E-DIII flexibility is needed for optimal virus infectivity.

**Figure 3:**
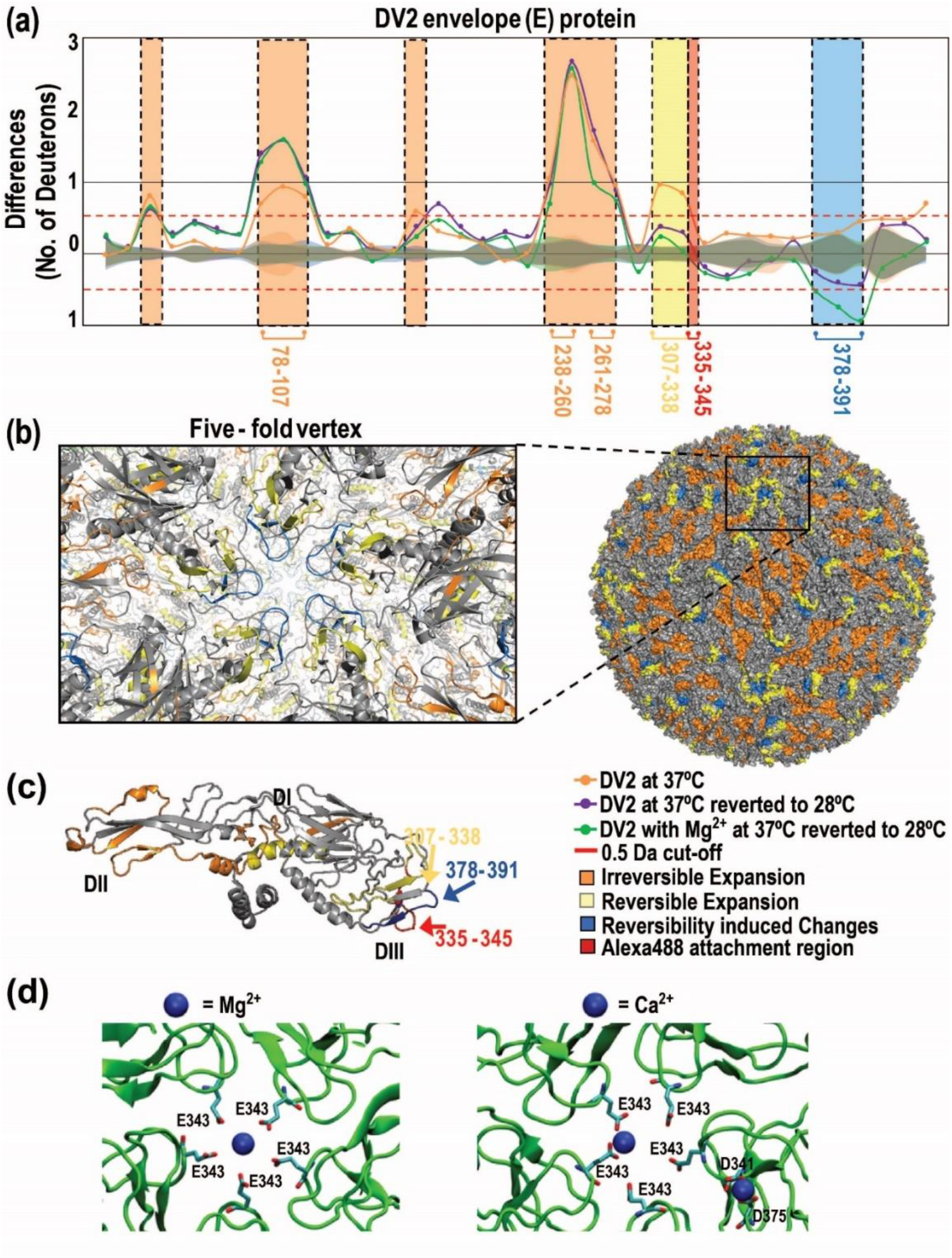
Non-uniform temperature-specific changes in DENV2 in absence and presence of 1 mM Mg^2^+ at 37 and 28°C by HDXMS and equilibrated coordination of the active Mg^2^+ or Ca^2^+ion binding sites. **(a)** HDXMS Difference plot for temperature-induced differences (t = 1 min) in E protein from DENV2 at 37°C (orange line) and after reversal of temperature to 28°C in absence (violet line) and presence (green line) of 1 mM Mg^2^+-ions. The difference plot displays the differences in exchange between the two temperatures indicated in peptides protein-wide (Y-axis) with each dot representing a pepsin fragment peptide, listed from the N-to C-terminus. Y-axis-differences in deuterons, X-axis-pepsin fragment peptides. Differences in deuterium exchange above 0.5 D are considered significant (red dash line). Standard error for each peptide is shown as overlapping shaded regions along the X-axis and colored according to the conditions in the difference plots. Orthogonal views of the differences in deuterium exchange in **(b)** viral particle surface and (**inset**) five-fold vertex and **(c)** E protein peptides with temperature changes mapped onto the respective Cryo-EM structures (PDB ID: 3J27). Regions with yellow color represents regions (V_307_–K_338_ & G_426_–A_445_) with reversible expansion, orange color represents regions (G_78_–F_107_ & T_238_–F_278_) with irreversible expansion and blue color represents regions (I_378_–F_391_) with reversibility induced changes, when virus temperature was lowered to 28°C. Red color indicates the Alexa488-TFP-attachment site (I_335_–V_345_) on E protein. **(d)** Snapshots of the E protein showing pentamer interface showing number of atomic contacts within 3.0 nm between Mg^2^+/Ca^2^+ cations and the I_335_–V_345_ peptide.

Additionally, peptide 378_I_-391_F_ constitutes the apex region of the junction where E protein units are arranged into five-fold vertices (represented by blue color; Figure 3b and Inset 3b), indicating that the divalent cation dependent E-DIII conformational changes belong to the DENV2 five-fold vertices. Taken together, our data identify the DENV2 E protein regions that undergo Mg^2^+ and Ca^2^+ ion-dependent reversible large scale conformational changes. Furthermore, both in absence (Figure 2b and S2e) and presence (Figure 2c and S2f) of divalent cations, no or moderate changes were observed for DENV 1, consistent with evidence from literature that DENV 1 does not show large scale morphological changes in this temperature range (22, 23).

### Divalent cation binding disrupts the inter-subunit salt bridges at 5-fold vertex of DENV2

We investigated the interaction between Mg^2+^/Ca^2+^ ions and the surface E proteins at the 5-fold symmetry vertices of the “bumpy” DENV2 virus, using explicitly solvated, atomic-resolution molecular dynamics (MD) simulations. Overall, MD simulations suggested that Mg^2^+ ion interacts with the Glu 343 (E343) while Ca^2^+ ion interacts either with Glu 343 or two Asp residues 341 (D341) and 375 (D375) (Figure 3d). Interestingly, E343 and D341 lie within the Alexa488-TFP attachment region spanning peptide 335_I_-345 (Figure 3c) of the DENV2 E-DIII. Additionally, residue 375 is in close proximity to the region showing Mg^2^+ion dependent reversible large scale conformational changes (Figure 3c). Again, E343 and D341/D375 lie within peptides that constitute the apex region of the junction where E protein units are arranged into five-fold vertices (represented by blue color; Figure 3b and Inset 3b), supporting the notion that divalent-ions bind at DENV2 five-fold vertices.

Furthermore, the initial simulation snapshots of the pentameric interface showed spontaneous formation of multiple inter-subunit salt bridges between E343 and K344 from adjacent subunits i.e. between chains I-II, II-III, IV-V and V-I (Figure 4a). These salt bridges presumably stabilize/lock the protein-protein interfaces in the “bumpy” state of the virion. The electrostatic potential and visualization of the resultant field lines of the E protein pentamer (Figure 4b and 4c) revealed the presence of an electrostatic “funnel” expected to drive divalent cations towards the 5-fold vertex of the symmetry, thus increasing the local divalent cation concentration. Indeed, after 200 ns of simulation, Ca^2^+ ions were found to have stably associated with the E343 side chain carboxylate oxygen of three (chains: I, IV and V) out of five chains (Figure 4a). Comparisons between the initial and the final snapshots of pairs of E protein interfaces and the corresponding time-series plots for chains I-II (Figure 4d) showed that the salt bridge remained stable over the course of the simulation where Ca^2^+ ion did not bind to E343. On the other hand, a concomitant breakage of the E343-K344 salt bridge was observed in the case of chains IV-V (Figure 4e) and V-I (Figure 4f) due to binding of Ca^2^+ ion to E343 at ~20 ns and ~120 ns, respectively.

**Figure 4.**
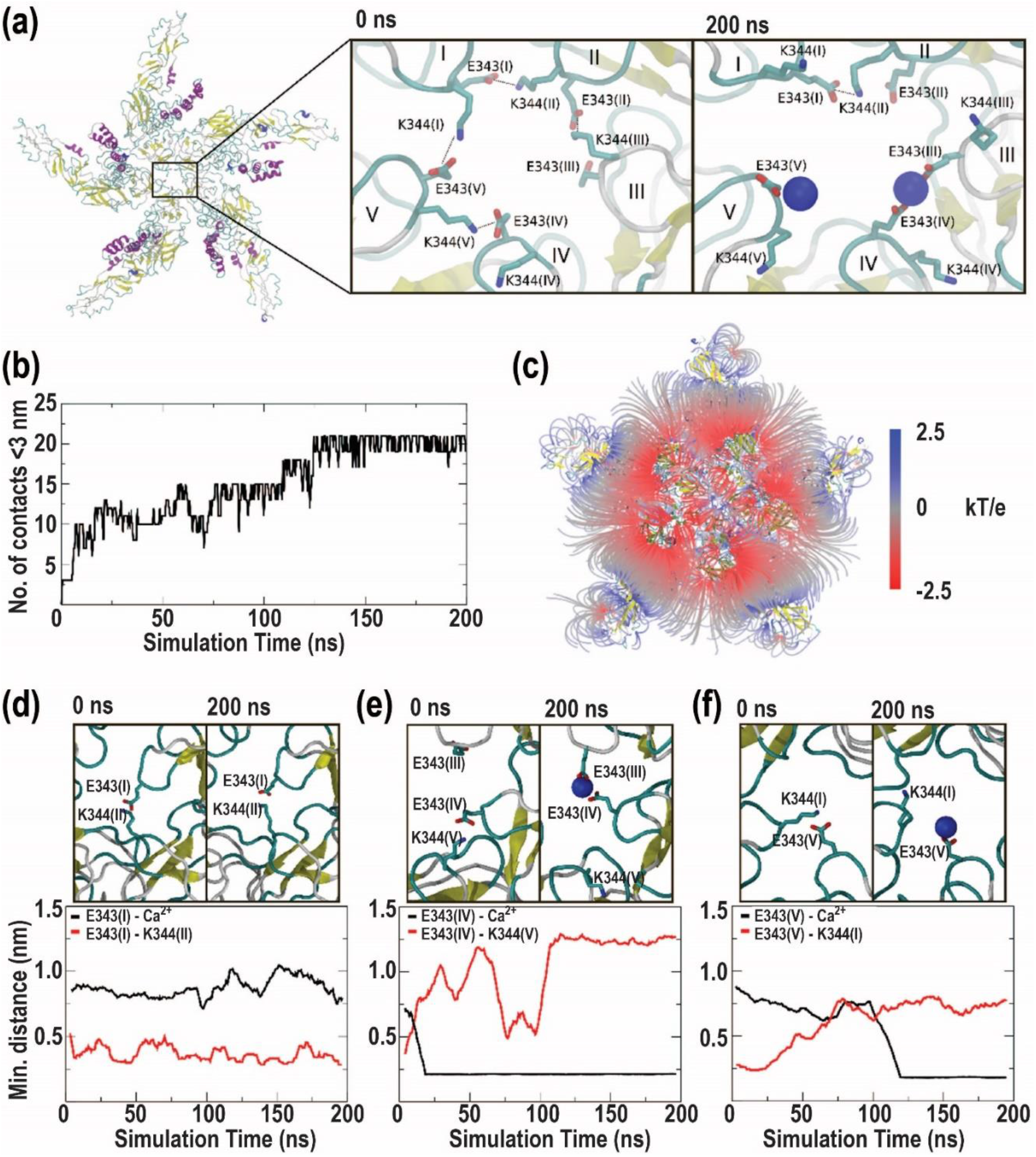
Molecular dynamics simulations reveal an electrostatic funnel for calcium binding and salt bridge disruption at the E protein 5-fold vertex. **(a)** Initial and final snapshots of the pentamer (bottom view). **(b)** Number of atomic contacts within 3.0 nm between Ca^2^+ cations and the 335_i_–345_V_ peptide. **(c)** Lines indicate the direction of the electrostatic field for the E protein pentamer (top view), colored by potential energy value from -2.5 kT/e (red) to 2.5 kT/e (blue). **(d)**, **(e)**, and **(f)** show initial and final snapshots of the interface between chains I-II, IV-V, V-I, respectively. Below each snapshot, corresponding time-dependent minimum distances are shown for E343-Ca^2^+ and E343-K344. In all protein snapshots, the secondary structure is shown in cartoons representation, with residue side chains in CPK wireframe format, and Ca^2^+ cations shown as blue spheres.

Based on these observations, we hypothesize that divalent cations have the ability to “soak” the protein 5-fold interface, breaking the inter-chain salt bridges by binding to acidic residues, and, thereby, reducing the stability of the “bumpy” state. Moreover, this notion needs to be supported by mutational studies, but such experiments are beyond the scope of the present study.

### DENV2 E-DIII loses temperature and divalent cation dependent flexibility at 40°C

At later stages of dengue infection, during dengue fever, DENV is further exposed to temperatures as high as 40°C. Thus, we investigated the temperature dependent E-DIII large scale conformational changes when DENV2 and DENV1 transit to 40°C before reverting to 25°C.

An increase in temperature from 37°C to 40°C, resulted in no or marginal increase in the values of <*τ*_f_> (Figure 5a) and α1 (Figure S4a) for DENV2, indicating no further DENV2 E protein conformational changes, in line with literature (22). With the stepwise reversal in temperature from 40°C to 25°C in steps of 4°C, marginal or no change in the values of <*τ*_f_>(Figure 5) and α1 (Figure S4) were observed both in the absence and the presence of divalent cations. This absence of DENV2 E protein large scale conformational changes contrasts with the effects observed between 25°C and 37°C in the presence of Mg^2+^ and Ca^2+^ ions, indicating irreversible bumpy morphology of DENV2 at 40°C. Thus, in absence of any further large scale conformational changes, as compared to that of 37°C (22), the formation of an irreversible bumpy morphology of DENV2 at 40°C probably indicates the faulty binding of divalent cations which hindered their ability to “soak” the E protein at 5-fold interface. Interestingly, recent study with Zika virus supported that hydrogen-bonding interaction network within the ZIKV envelope protein contribute to the virus structural stability as well as in vivo pathogenesis (52).

**Figure 5:**
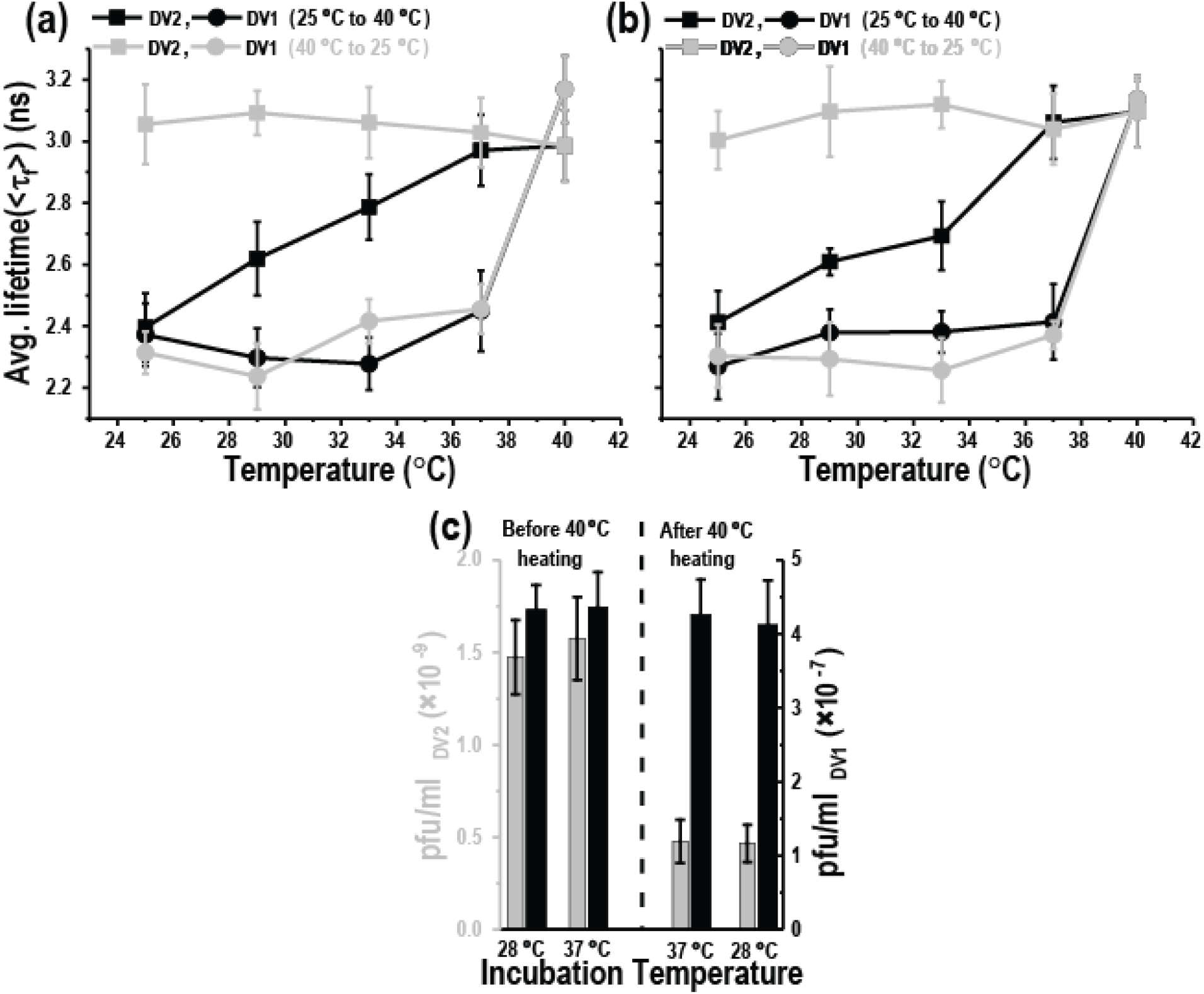
Structural transitions and infectivity of DENV2 and DENV1 between 25°C and 40°C in (a) absence and (b) presence of Mg^2^+-ions. **(a)** The average fluorescence lifetimes (<τ_f_>) for dual-labeled 5 × 10^7^ pfu/ml DENV2 (squares) and 5 × 10^7^ pfu/ml DENV1 (circles). The data was collected during temperature increase from 25°C to 40°C (black circles/squares) followed by decrease from 40°C to 25°C (grey circles/squares). **(b)** The average fluorescence lifetimes (<τ_f_>) for dual-labeled DENV2 (squares) and DENV1 (circles) in the presence of 1 mM Mg^2^+-ions. The data was collected during temperature increase from 25°C to 40°C (black circles/squares) followed by decrease from 40°C to 25°C (grey circles/squares). Error bars represents standard deviations of six different experimental replicates in both DENV2 and DENV1 viruses. **(c)** Histogram showing Plaque forming virus titers performed at 1 hour incubation of either DENV2 (grey bars) or DENV1 (black bars) with BHK21 cells at temperatures of either 28°C or 37°C. Plaque assay was performed with both viruses before and after experiencing 40°C to have attained either flexible or locked E protein conformation. Error bars represents standard deviations of three different experimental duplicates in both DENV2 and DENV1 viruses.

Furthermore, both, in absence (Figure 5a and S4c) and presence (Figure 5b and S4d) of divalent cations, an increase in temperature from 37°C to 40°C, resulted in increased values of <*τ*_f_> (2.4 ± 0.05 ns → 3.2 ± 0.05 ns) (Figure 2b) and α1 (43 ±3 % → 65 ± 4 %) (Figure 2a) for DENV1. Again, the difference in the values of α1 at 37°C and 40°C indicated that, similar to DENV2, only ~22% of DENV1 underwent large scale morphological changes. In contrast to DENV2, these DENV1 E protein large scale conformational changes were reversible when reverting to temperatures below 37°C. The results showed that DENV1 exhibits reversible large scale morphological changes in the temperature range from 25°C to 40°C. In addition, these changes are independent of the presence of divalent cations and thus may indicate a higher structural stability of DENV1, due to stronger intra-dimeric E protein contacts, as compared to DENV2(21).

### DENV2 loses virus infectivity due to loss of divalent cation dependent E-DIII flexibility at 40°C

Since, the smooth and the bumpy form of DENV2 do not differ in their infectivity (20), we monitored the effect of DENV2 E-DIII large scale conformational flexibility by measuring infectivity on mammalian BHK21 cells. For this, DENV2 were incubated at 25°C, 37°C and 40°C to attain the smooth, the partial reversible bumpy and the irreversible bumpy morphologies, respectively. Interestingly, the number of plaques produced by the irreversible bumpy DENV2 particles (virus incubated at 40°C) was found to be ~3-fold smaller than the number of plaques generated either by the partially reversible bumpy DENV2 (virus incubated at 37°C) or the smooth DENV2 (virus incubated at 25°C) (Figure 5c, grey bars). The results are in line with the involvement of E-DIII in the host receptor binding process (42, 43, 50, 51). The results also indicate that DENV2 E-DIII large scale conformational flexibility probably plays a role during infectivity of both the smooth and partial bumpy form of DENV2. Moreover, loss of such E-DIII flexibility probably inhibits the interaction of E-DIII with host receptors in the irreversible bumpy DENV2 morphology.

We supported the DENV2 infectivity loss by using DENV2-specific human monoclonal antibody (HMAb) 2D22, which is therapeutic in a mouse model of antibody-enhanced severe dengue disease (53). HMAb 2D22 binds across DENV2 E proteins in the dimeric structure, and locks E-DIII large scale conformational changes both before and/or after DENV2 morphological changes at 37°C (19). Thus, HMAb 2D22 bound DENV2 at 37°C should mimic the irreversible bumpy morphology of DENV2, formed due to heating at 40°C. Thus, no changes in DENV2 E protein conformations should be observed either in absence or in presence of divalent cations.

To achieve this, HMAb 2D22 was added to either the smooth (at 25°C) or bumpy (at 37°C) form of DENV2, in presence of either Mg^2^+or Ca^2^+ions. As expected, no significant changes in the values of the <*τ*_f_> and α1 (from mean value of 2.2 ± 0.1 ns and 50 ± 4 %, respectively) were recorded with the stepwise increase in temperature from 25°C to 37°C (Figure S5a, S5b and S5c), suggesting that HMAb 2D22 can bind to and lock the E-DIII domain in smooth DENV2. Similarly, addition of HMAb 2D22 to DENV2 at 37°C caused no decrease in the values of the <*τ*_f_> and α1 (from mean value of 3.0 ± 0.1 ns and 72 ± 3 %, respectively), with the stepwise reversal of temperature from 37°C to 25°C (Figure S5a, S5b and S5c). This result indicates that HMAb 2D22 can also bind to and lock the E-DIII domain in bumpy morphology of DENV2. Taken together our results validated that HMAb 2D22 exerted loss of E-DIII large scale conformational flexibility, in both the smooth and the bumpy form of DENV2, and results in a reduction of viral infectivity (53, 54). By analogy, loss of E-DIII large scale conformational flexibility due to exposure to temperature of 40°C or above may result in the reduction of DENV2 infectivity (Figure 5c, grey bars).

Since, infectivity of both the smooth (at 25°C) and the bumpy (at 37°C and 40°C) DENV2 morphologies get affected due to the loss of DENV2 E-DIII large scale conformational flexibility, this indicates a limited role of the specific DENV morphologies. This notion is further strengthened by the DENV1 infectivity results. DENV1, like DENV2, shows large scale morphological expansion but unlike DENV2, does not loose E protein conformational flexibility after experiencing 40°C. In addition, we observed no or marginal difference in the number of plaques produced (Figure 5c, black bars) by DENV1 in both, either before or after experiencing 40°C, conditions. However, we are monitoring the distance changes only between DENV2 E-DIII or DENV1 E-DIII/E-DII to their respective viral bilayers and thus, cannot exclude the possibility that the overall E protein conformations could be different in presence of HMAb as compared to that of at 40°C.

Although, the loss of temperature dependent DENV2 E-DIII large scale conformational flexibility is clearly distinguishable at 25°C, the differences between flexible and non-flexible E-DIII either at 37°C or at 40°C are indistinguishable (Figure 5a and 5b). Taken into account the similar DENV2 bumpy morphology both at 37°C and at 40°C (22), the ~3-folds difference in DENV2 infectivity cannot be explained (Figure 5c) on the basis of the E protein large scale conformational changes. On the other hand, it is equally intriguing that DENV2 infectivity does not change between 25°C and 37°C despite strong temperature dependent E protein large scale conformational changes (15, 22, 23) (compare grey bars in panel marked as “before 40°C heating” in Figure 5c). This implies that large scale E protein conformational changes are not the basis of DENV infectivity.

### DENV2 shows reduced E-DIII dynamics at 40°C as compared to the range between 25°C and 37°C

Since, the large scale E-DIII conformational changes that are temperature dependent cannot be the basis of DENV infectivity, we investigated what role DENV intrinsic dynamics at faster time and smaller length scales play in viral infectivity. Thus, we investigated the E protein intrinsic dynamics w.r.t the viral bilayer for discrete DENV morphologies that lie between the smooth and bumpy forms of the virus.

To achieve this, we used FRET-FCS that analyses the fluctuations in FRET efficiency caused by protein conformational fluctuations. As the fluorescence intensities change when a particle moves through a laser focus, FRET-FCS measures the so-called proximity ratio *(p*), which is a function related to the FRET efficiency (Explained in Figures S6a) (55, 56) and which depends strongly on the separation between donor and acceptor but not on the position of the molecule in the observation volume. Therefore, the correlation function of the proximity ratio *p (Gp)* is expected to have contributions mainly from structural dynamics, and minimally from molecular diffusion. In addition, by constructing the *Gp*, rather than of the fluorescence intensity, we simplify the extraction of intramolecular kinetics or intrinsic dynamics for the correlation function. The approach was calibrated (see Figure S6) and used for DENV measurements. Fitting of*Gp* with stretched exponential provides an effective relaxation time (*τ_p_*) associated with the correlated motion at smaller length scales, and the stretch parameter (β) describing the heterogeneity of the system. β can vary between 1 (where the system displays normal two-state Arrhenius kinetics, with one discrete energy barrier) and 0 (where there is a continuum of equal energy barriers and the system shows power-law kinetics).

Next, a stepwise increase in temperature from 25°C to 37°C, either in presence or in absence of Mg^2^+ions, resulted in no or marginal increase in values of *τ_p_* (~0.9 ms) (Figure 6c and 6d) and β (~0.5) (Figure S6c and S6d) for both DENV2 and DENV1. The absence of any changes in intrinsic dynamics is more surprising for DENV2 as it undergoes changes from the smooth to the bumpy form between these two temperatures.

Fitting of DENV2 *Gp* (Figure 6a (blue curve)) at 25°C in presence of Mg^2^+ions, yielded values for *τ_p_* and β of 0.9 ± 0.1 ms (Figure 6c) and 0.48 ± 0.06 (Figure S6c), respectively. In addition, the value obtained for β confirms the DENV2 heterogeneity and existence of more than two E protein conformational states. Moreover, similar values for *τ_p_* = 0.89 ± 0.1 ms (Figure 6d) and β = 0.5 ± 0.04 (Figure S6d) were obtained for DENV2 E-DIII intrinsic dynamics in absence of Mg+^2^ions, suggesting the limited role of divalent cations for protein dynamics in contrast to the morphological changes. Next, fitting of DENV1 *Gp* (Figure 6b) provided similar values for *τ_p_* (0.9 ± 0.1 ms and 0.9 ± 0.1 ms in presence and absence of Mg^2^+ions, respectively) and β (0.51 ± 0.04 and 0.51 ± 0.04 in presence and absence of Mg^2^+ions, respectively) as for DENV2. The similarities in *τ_p_* and β values between DENV2 and DENV1, suggests that both DENV E proteins shows similar intrinsic dynamics at smaller length scales.

**Figure 6:**
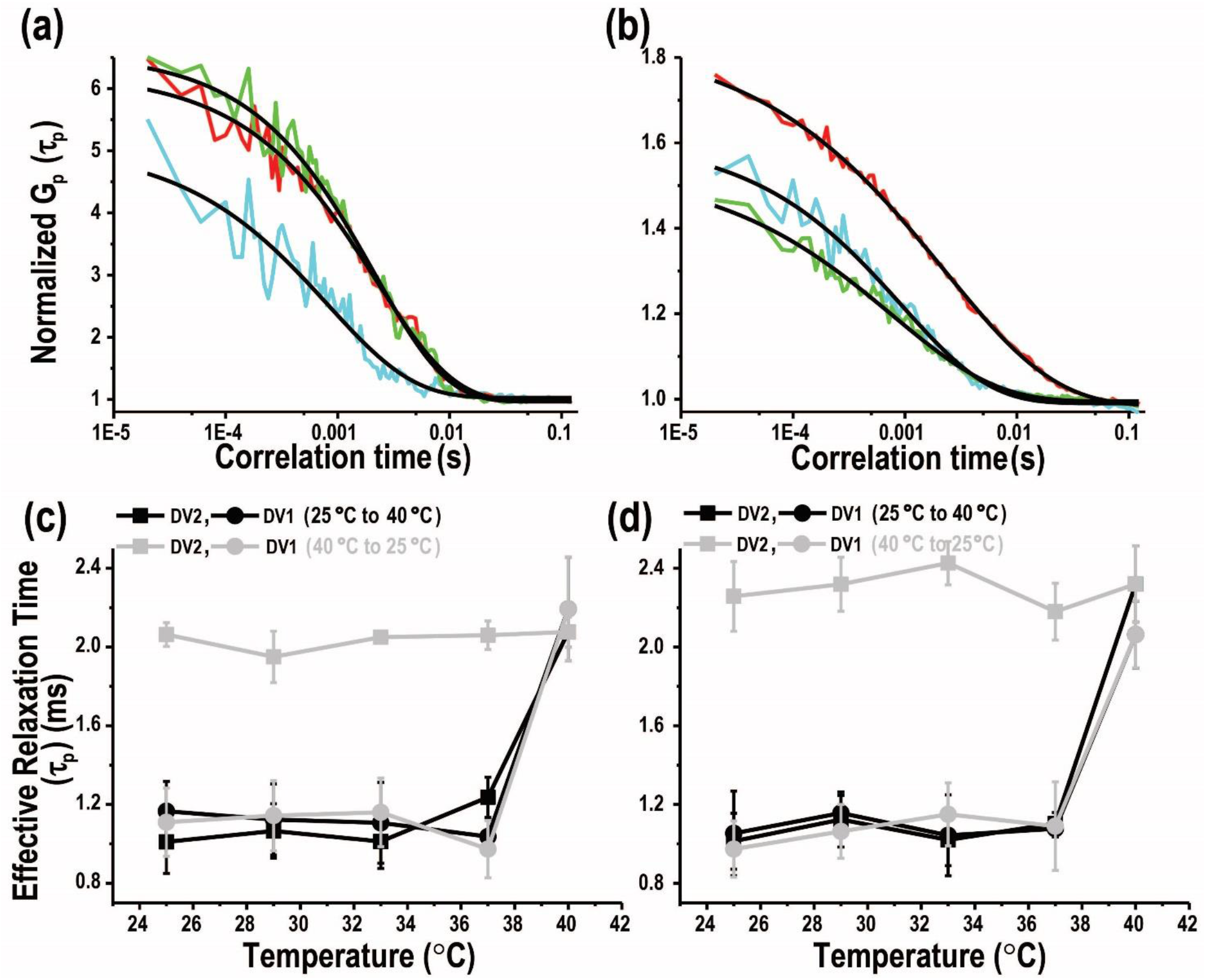
Fluctuation dynamics of DENV2 and DENV1 between 25°C and 40°C in absence and presence of Mg^2^+-ion. Autocorrelation curves of proximity ratio for 5 × 10^7^ pfu/ml **(a)** DENV2 and 5 × 10^7^ pfu/ml **(b)** DENV1 in presence of 1 mM Mg^2^+-ions. Representative curves are plotted at 25°C (cyan traces), after temperature increase from 25°C until 40°C (grey traces) and finally followed by decrease from 40°C to 25°C (green traces). The black line represents the best fit to stretched exponential and the obtained values are discussed in the article. Effective relaxation time, τ_P_, calculated by fitting these autocorrelation curves of proximity ratio for DENV2 (squares) and DENV1 (circles) either in **(c)** presence or in **(d)** absence of 1 mM Mg^2^+-ions. The data was collected during temperature increase from 25°C to 40°C (black circles/squares) followed by decrease from 40°C to 25°C (grey circles/squares). Error bars represents standard deviations of four different experimental replicates in both DENV2 and DENV1 viruses.

Furthermore, an increase in temperature from 37°C to 40°C resulted in an ~2.5-fold increase of *τ_p_* for both DENV2 (2.1 ± 0.05_+Mg_ and 2.3 ± 0.1_-Mg_) and DENV1 (2.1 ± 0.05_+Mg_ and 2.3 ± 0.1_-Mg_) (Figure 6d), indicating the adverse effect of dengue fever conditions on DENV E protein intrinsic dynamics.

Interestingly, no change in the values of *τ_p_* (Figure 6c and 6d) was observed for DENV2, with the reversal in temperature from 40°C to 25°C. The result reveals the correlation between the losses of E protein flexibility, at high temperature, and the decrease in its intrinsic dynamics. Consequently, this irreversible ~2.5-fold reduction in DENV2 E-protein intrinsic dynamics in the non-flexible conformation at ≤37°C, may be responsible for decreased DENV2 infectivity (Figure 5c). Similarly, lack of change in DENV2 intrinsic dynamics at 25°C and 37°C (before experiencing 40°C) coincides well with the unchanged DENV2 infectivity either in the smooth or the bumpy morphology (20) (Figure 5c). Thus, the results indicate the role of virus intrinsic dynamics as compared to virus morphologies.

This notion was further supported by the reversibility of *τ_p_* (~2.1 ms → ~0.9 ms) for DENV1 (both in presence (Figure 6c) and absence (Figure 6d) of divalent cations from 40°C → 25°C) that correlates well with unperturbed DENV1 infectivity, even after exposure to 40°C (Figure 5c, black bars). The results indicate that the reversible E protein intrinsic dynamics (~0.9 ms → ~2.1 ms → ~0.9 ms) may lead to reversed large scale conformational changes as seen with DENV1 between 37°C and 40°C. Similarly, no change in this E protein intrinsic dynamics over temperature range (25°C to 37°C) correlates with unperturbed DENV2 infectivity in both smooth and bumpy forms (Figure 5c). Therefore, DENV E protein intrinsic dynamics but not its static conformations are correlated to viral infectivity and explain the differences in DENV1 and DENV2 infectivity.

## Summary

Various Dengue virus strains undergo morphological changes from a smooth low temperature morphology to a bumpy morphology above 35°C. This transition called “breathing” (14, 15, 57, 58) is similar to the “breathing” effect observed in icosahedral picornaviruses that reversibly expose cleavage sites to proteases (17, 46–49). Thus, by analogy, DENV “breathing” implies some reversible conformational changes of DENV envelope proteins. However, earlier studies showed that heating the smooth low temperature DENV2 virus to temperatures above 35°C produced an irreversible bumpy morphology that did not change even when reducing the temperature below 35°C (19, 20, 22, 23). These studies, though, were conducted in the absence of divalent cations which could be the reason for the persistence of irreversible viral transitions that would prevent DENV E protein to undergo picornavirus-like reversible large scale conformational changes. Here, therefore, we demonstrated that the addition of Mg^2^+ or Ca^2^+ ions at physiological concentrations induces a partial reversibility, underlining the importance of divalent cations for viral functionality. However, it should be noted that we observed large scale conformational changes in only ~15-20% of DENV. This structural heterogeneity arising due incomplete cleavages of precursor-to-membrane (prM) or due to other perturbations, helps viruses to avoid neutralization by antibodies (13).

Nevertheless, we were not able to correlate the envelope conformations to infectivity. DENV E protein large scale conformational changes can be inhibited by using antibodies that can lock the virus in both, the smooth and the bumpy morphologies with reduced infectivity. However, unlike the antibody locked irreversible morphology, temperature induced irreversible bumpy DENV2 morphology does not reduce viral infectivity in BHK21 cells (20). Furthermore, plaque assays demonstrated that the conformational state of the virus envelope of DENV1 and DENV2 are not correlated to viral infectivity.

We therefore investigated the E protein intrinsic dynamics at the smaller distances and faster time scales, for both smooth and bumpy form, under the same conditions. The intrinsic dynamics measured by FRET-FCS indicated that for DENV1 E protein the intrinsic dynamics as well as infectivity did not change between 25°C to 40°C. DENV2 E protein, however, showed a 2.5-fold decrease in intrinsic dynamics at 40°C compared to the dynamics between 25°C -37°C, and also showed a concomitant decrease in infectivity (Figure 7).

**Figure 7:**
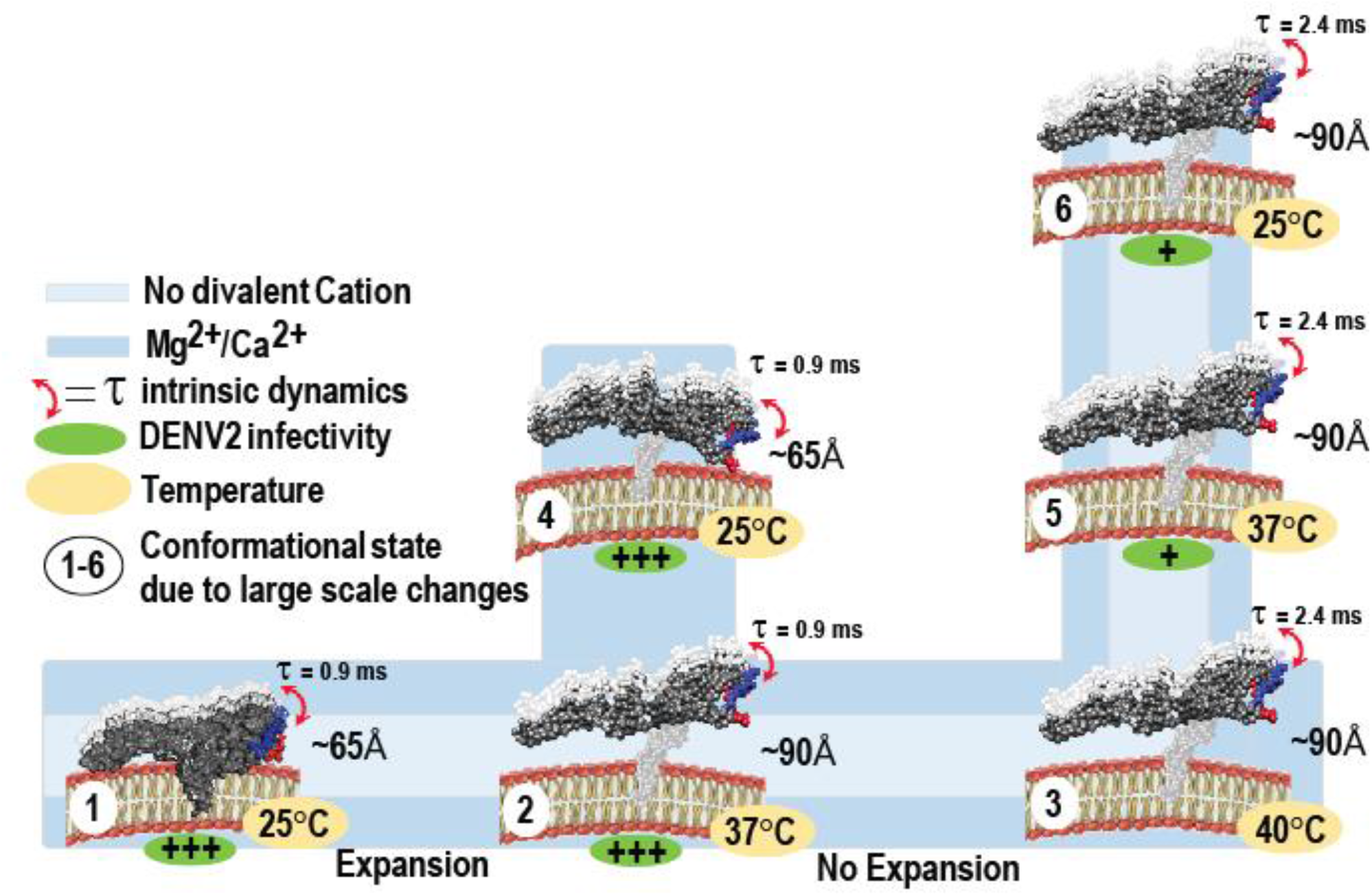
Proposed model differentiating type of DENV2 “breathing” and its relation to virus infectivity. The diagram summarizes divalent cation and temperature dependent large and small scale changes in DENV2 E-DIII at five-fold vertex. (1) In smooth virus (PDB ID: 3J27) E-DIII is buried in pentameric units at an average distance of ~65 Å from the viral bilayer and showed intrinsic dynamics in ~0.9 ms range. During viral expansion (PDB ID: 3ZKO), both in presence (blue color) and absence (light blue color) of divalent cations, upto either (2) 37°C or (3) 40°C, E-DIII undergo large scale conformational changes and the average distance increases to ~90 Å from 65 Å, but showed intrinsic dynamics in ~0.9 ms and ~2.4 ms range at 37°C and 40°C, respectively. (4) E-DIII undergo temperature dependent large scale conformational contraction only in the presence of divalent cations, with lowering of temperature from 37°C to 25°C. The average distance between E-DIII and bilayer decreased from ~90 Å to 65 Å. When heated to 40°C and lowering of temperature either to (5) 37°C or (6) 25°C, E-DIII loses its ability to undergo large scale temperature dependent conformational changes and thus, no change in either distance between E-DIII w.r.t bilayer or domain intrinsic dynamics were observed. Under such conditions, DENV2 showed reduced infectivity (+ve sign in green oval) in BHK21 cells.

This work on the large scale conformational changes and intrinsic dynamics of DENV E proteins shows a remarkable adaptation of the virus to its two hosts’ temperatures to attain infectivity. It shows not only the importance of divalent cations for the structural and dynamic investigation of viruses but also demonstrates that dynamics complements the DENV structural studies and is critical for determining factors influencing DENV infectivity. These dynamic events are essential aspects of the DENV life cycle and can be targeted for antiviral strategies. Lastly, we speculate that mutations targeting DENV intrinsic dynamics instead of large scale conformational changes may be instrumental in producing attenuated viruses that are capable of invoking neutralizing antibodies.

## Methods

### Dyes and Viruses

Both alexafluor 488 TFP ester (AF488-TFP) and 1,1‘-Dioctadecyl-3,3,3’,3’-Tetramethylindocarbocyanine Perchlorate (DiI-C18) were purchased from Invitrogen (Singapore). DENV2 NGC and DENV1 PVP159 were produced and purified as previously described (20, 22).

### Virus labeling

Alexa Fluor 488 TFP ester (AF488-TFP) and 1,1‘-Dioctadecyl-3,3,3’,3’-Tetramethylindocarbocyanine Perchlorate (DiI-C18) were purchased from ThermoFisher Scientific, Singapore. Purified viruses had stock concentrations of 6 × 10^10^ plaque forming units (PFU)/ml for DENV2 (NGC) and 4.2 × 10^10^PFU/ml for DENV1 (PVP 159). Both DENV1 (PVP 159) and DENV2 (NGC) were dual labelled by sequential addition of DiI-C18 and AF488-TFP, in buffer containing 10 mM HEPES, 150 mM NaCl at pH 7.4 (HN buffer). The DiI-C18 and AF488-TFP stock solutions were prepared in dimethylsulfoxide and subsequently their concentrations were determined using molar extinction coefficients of 144,000 M^-1^cm^-1^ and 71,000M^-1^cm^-1^, respectively. Initially, the DiI-C18 stock solution was diluted in 50 μl of HN buffer to a final concentration of 50 nM and sonicated for 10 min. Next, ~2.5 × 10^8^ PFU of the purified viruses were directly added to the DiI-C18 containing HN buffer solution and incubated for 1h at 4°C to label the virus-lipid bilayer. Subsequently, AF488-TFP at a final concentration of 750nM was added to the DiI-C18 labelled virus HN buffer mixture and incubated for an additional hour at room temperature. The free dye molecules were removed by gel filtration (MicroSpinTM S-200 HR columns, GE Healthcare, Singapore). As a control, single labelled donor-only viruses were prepared by labelling with AF488-TFP only.

### Plaque-forming assay

BHK cells were cultured in RPMI supplemented with 10% FBS at 37°C in 5% CO2. When cells reached approximately 80% confluence, virus infection was carried out. Virus was serially diluted 10-fold from 1:10^2^ to 1:10^10^ using RPMI supplemented with 10% FBS. For the experiment, the virus was preincubated at 37°C or 40°C for 30 min and then 100 μl of virus was layered onto the BHK cells, which were preequilibrated at 37°C for 2 h. The cells were then further incubated at their respective temperatures for 2 h. The infected cells were overlaid with RPMI supplemented with 10% FBS and 1% Aquacide. The plates were further incubated at 37°C in 5% CO2 for 6 days. The cells were then stained by first discarding the supernatant and then adding 1 ml of 0.5% (wt/vol) crystal violet-25% formaldehyde to the cell monolayer and further incubated for 1 h. The cell monolayers were then washed with water, and plaques were counted by visual inspection. The experiments were repeated at least thrice.

### Time resolved Förster Resonance Energy Transfer (trFRET) measurements

tr-FRET measurements were carried out on a commercial Olympus FV1200 laser scanning confocal microscope (IX83, Olympus, Singapore) equipped with a time-resolved LSM upgrade kit (Microtime 200, PicoQuant, GmbH, Berlin, Germany). The samples were excited with a 485 nm pulsed diode laser with a 20 MHz repetition rate and 29 μW power (PDL series, Sepia II combiner module). The beam was focused into the sample by a water immersion objective (× 60, NA 1.2; Olympus, Tokyo, Japan) after being reflected by a dichroic mirror (DM405/485/543/635 band pass, Olympus, Singapore) and the scanning unit. The fluorescence was collected by the same objective followed by a pinhole (120 μm) to remove out-of-focus light. The fluorescence signal was spectrally divided into donor (green) and acceptor (red) channels by a 560DCLP dichroic mirror. The donor fluorescence was recorded by a set of single molecule avalanche photodiodes (SPADs) (SPCM-AQR-14, PerkinElmer Optoelectronics, Quebec, Canada), through a 520/35 band pass emission filter (Omega, VT). This donor signal was further processed by a time correlated single photon counting card (TimeHarp 260, PicoQuant) to build up the histogram of photon arrival times.

The temperature of the sample was controlled by an on-stage incubator (TempControl 37-2 digital, Pecon, Erbach, Germany) and an objective heater (TC-124A, Warner Instruments, Hamden, CT). The TR-FRET measurements were recorded for 180 s after incubating single/dual-labelled virus samples for at least 30min, at a given temperature. The mean lifetime angle (τ) was calculated from the individual fluorescence lifetimes (τi) and their relative amplitudes (αi) according to (τ)=Σαiτi. Donor fluorescence lifetime decay data were treated using the software SymPhoTime 64 (PicoQuant, GmbH). In all cases, the χ^2^ values were close to 1 and the weighted residuals as well as their autocorrelation were distributed randomly around 0, indicating a good fit. The reported values are mean and s.d.’s from six replicates for both DENV1 (PVP 159) and DENV2 (NGC) from 3 different purified virus batches.

### Identification of Biotin-labelled sites on DENV2 proteins

The C, E and M-proteins from biotin-labelled DENV1 and 2 particles were separated by SDS PAGE analysis. The protein bands were excised from the SDS-PAGE gel and cut into fine bits after fixation with 50% Methanol containing 12% Acetic acid for 30 min at room temperature. Finely-cut gels were washed with 50 mM NH4HCO3 in 50% acetonitrile and dehydrated with 100% acetonitrile. Cysteines in the sample were reduced by treatment with 5 mM TCEP (Tris (2-carboxyethyl) phosphine) in 100 mM NH4HCO3 for 60 min at 57 °C and alkylated with 10 mM MMTS in 100 mM NH4HCO3. The protein bands were digested with trypsin (12.5 ng/μL trypsin in 500 mM TEAB (Triethylammonium bicarbonate)) overnight at 37 °C. Trypsin fragmentation peptides were eluted from the gel bits using 50 mM TEAB and desalted using Sep-Pak^®^ tC18 μElution Plate (186002318, Waters Corp., Milford, MA). The peptides were lyophilized and resuspended in PBS buffer. The biotinylated peptides were affinity purified using immobilized Neutravidin beads (Thermo Fisher Scientific) and the bound biotinylated peptides were eluted with 0.2% trifluoro-acetic acid, 0.1% formic acid in 80% acetonitrile. Eluted peptides were lyophilized and resuspended in PBS buffer before mass spectrometric analysis. Peptide identification was by an online LC MS/MS using TripleTOF 5600 system (AB SCIEX, Foster City, CA) in Information Dependent Acquisition Mode. The raw mass spectra were identified by searching against the DENV structural proteins database containing amino acid sequences of the C-, E-and M-proteins using the ProteinPilot 4.5 software (July 2012, AB SCIEX) that included variable modification (size-226.0776 Da) on lysine residues.

### Reversibility of DENV2 expansion in the presence and absence of Mg^2^+ by HDXMS

Purified DENV2 particles (0.25 mg/mL of E-protein) in NTE buffer were incubated at 28°C and 37°C for 30 min as described(22). In order to assess the reversibility of DENV2 expansion, DENV2 particles were first incubated at 28°C for 30 min and expanded by incubation at 37°C for 30 min followed by incubation at 28°C for 30 min. To test the reversibility of temperature-dependent DENV2 expansion in the presence of Mg^2^+ ions, the DENV2 particles that were pre-incubated in NTE buffer containing 1mM of Mg^2^+, were subjected to similar temperature perturbations.

### Amide hydrogen/deuterium exchange mass spectrometry (HDXMS)

The deuterium exchange of DENV2 under the various specified temperature and Mg^2^+ ions conditions were initiated by a 10-fold dilution in the respective buffers that were reconstituted with 99.9% D2O resulting in 89.9% final D2O concentration. Deuterium exchange under these various conditions were carried for 1 min.

All deuterium exchange reactions were quenched by lowering the pH read to 2.5 upon addition of prechilled NaOH in GnHCl and Tris(2-carboxyethyl) phosphine-hydrochloride (TCEP-HCl) to achieve a final concentration of 1.5 M GnHCl and 0.25 M TCEP-HCl. Quenched reactions were maintained at 4 °C on ice to minimize back exchange. Viral membrane phospholipids in the DENV2 complexes were removed by adding 0.1 mg of titanium dioxide (TiO2) (Sigma Aldrich, St. Louis, MO) to the quenched mixture and incubated for 1 min with mixing every 30 s. TiO2 in the samples were removed with a 0.22 μm filter (Merck Millipore, Darmstadt, Germany) after 1 min of centrifugation at 13,000 rpm. All deuterium exchange reactions were performed in triplicate, and the reported values for every peptide are an average of three independent reactions without correcting for back exchange.

### Pepsin fragment peptide identification

Peptides of C-, E-and M-protein from DENV2 NGC were identified by searching the mass spectra of the undeuterated samples of purified DENV2 NGC against DENV2 NGC structural protein database containing amino acid sequences of the C-, E- and M-proteins using PROTEIN LYNX GLOBAL SERVER version 3.0 (Waters, Milford, MA) software. Mass spectra of peptides from a single undeuterated DENV2 NGC samples with precursor ion mass tolerance of <10 ppm, and products per amino acids of at least 0.2 with a minimum intensity of 5000 for both precursor and product ions were selected. Five undeuterated samples were collected and the final peptide list generated from peptides identified independently in at least 2 of the 3 undeuterated samples.

### Measurements of deuterium uptake and calculation of deuterium exchange differences

Deuterium uptake of each peptide were measured using DYNAMX Ver. 2.0 software (Waters, Milford, MA) by subtracting the average mass centroid of the peptide after 1 min of deuterium exchange with the average mass centroid of the corresponding undeuterated peptide. Differences in deuterium exchange for all peptides in DENV2 under the various temperature and Mg^2^+ perturbations were determined by subtracting centroid masses of deuterated peptides between two experimental conditions. Deuterium exchange differences from DENV2 E-proteins and displayed from N to C-terminus in the difference plot. The standard deviations for deuterium uptake in all peptides were determined and a difference of 0.5 Da was used as the significance threshold for any differences in deuterium uptake across the two states compared and agrees with the observed standard errors measured in deuterated peptides.

### Molecular Dynamics Simulations

Only low-resolution cryo-EM data are available for the “bumpy” expanded state of DENV2, with stem/transmembrane regions of E and M proteins absent. Therefore, a targeted MD (TMD) approach was used to drive a complete, high-resolution structure of the mature “smooth” DENV2 virion (PDB ID: 3J27(23)) towards a structure (PDB ID: 3ZKO(20)) of the expanded virion. During TMD, steering forces were applied in the form of a harmonic potential that is dependent upon the gradient of the root mean squared deviation (RMSD) from the target structure^3^. The initial coordinates for mature DENV2, consisting of 180 E and M proteins embedded within a lipid bilayer, were taken from our previous work.^4^ The fully solvated and equilibrated virion system was treated in a coarse-grained (CG) MARTINI(38) representation. Subsequently, the E protein ectodomain structure was biased towards that of the expanded state during TMD.

Following TMD, a single pentamer surrounding a 5-fold vertex of the expanded virion was back-mapped to atomic-resolution, using geometric projection followed by sets of energy minimization. The protein transmembrane domains were embedded in a biologically relevant lipid bilayer(59) using the CHARMM-GUI(60). This composition corresponded to palmitoyloleoyl phosphatidylcholine (POPC), palmitoyloleoyl phosphatidylethanolamine (POPE), and palmitoyloleoyl phosphatidylserine (POPS) in a ratio of ~6:3:1(59). The system was placed in a cubic box of ~25×25’13 nm^3^ and solvated with ~180,000 explicit TIP3P^8^ water molecules. CHARMM36 parameters were used to treat the lipid(61) and protein(62). Ca^2^+ cations were added at a 0.1 M concentration, with additional Cl^-^ ions used to neutralize the overall system charge. The total system size corresponded to ~840,000 atoms. The energy of the system was minimized with 10,000 steps using the steepest descents algorithm with a 0.1 nm step size. Equilibration in the *NVT* ensemble was performed for 5 ns using position restraints on protein heavy atoms with a force constant of 1,000 kJ mol^-1^ nm^-2^. Multiple sets of subsequent simulations were then performed in the *NPT* ensemble with gradually decreasing force constants on protein heavy atoms, to prevent unrealistic deformation of the pentamer in the absence of the remaining virion assembly: i) 5 ns at 1,000 kJ mol^-1^ nm^-2^; ii) 5 ns at 800 kJ mol^-1^ nm^-2^; iii) 5 ns at 600 kJ mol^-1^ nm^-2^; iv) 5 ns at 400 kJ mol^-1^ nm^-2^; and v) 180 ns at 200 kJ mol^-1^ nm^-2^.

All simulations were run using GROMACS 5.1.4 package(63). Equations of motion were integrated through the Verlet leapfrog algorithm with a 2 fs time step, and bonds connected to hydrogens were constrained with the LINCS algorithm. The cutoff distance was 1.2 nm for the short-range neighbour list and for van der Waal’s interactions with a smooth switching function from 0.8 nm. The Particle Mesh Ewald method was applied for long-range electrostatic interactions with a 1.2 nm real space cutoff.^12^ The Nose-Hoover thermostat^13,14>^ and Parinello-Rahman barostat^15^ were used to maintain the temperature and pressure at 310 K and 1 bar, respectively. Simulations were performed on an in-house Linux cluster, composed of nodes containing 2 GPUs (Nvidia K20) and 20 CPUs (Intel® Xeon® CPU E5-2680 v2 @ 2.8 GHz) each. Time dependent analyses were performed using GROMACS 5.1.4. Visualization of simulation snapshots used the VMD package.^16^ Electrostatic potential calculations were performed using the APBS plugin^17^ for the Chimera^18^ visualization package. Partial charges were obtained from the AMBER99SB force field. Residues were protonated according to their dominant populations at pH 7.0.

### FRET fluctuation correlation spectroscopy measurements (FRET-FCS)

A 40-base oligonucleotide 5’-GGGTT-(A)30-AACCC-3’ DNA hairpin-loop was purchased from IDT Technologies (Singapore). Donor fluorophore carboxytetramethylrhodamine (TMR) is attached at its 3’ end via a modified cytosine and a six-carbon linker. Acceptor fluorophore indodicarbocyanine (Cy5) is attached at its 5’ end via a three-carbon linker. The donor and acceptor form a fluorescence resonance energy transfer pair with a FRET distance (R_0_) of ≈5.3 nm. The structure of the fully closed hairpin-loop is illustrated in (Figure S5b).

FRET-FCS experiments were performed on same already mentioned commercial Olympus FV1200 laser scanning confocal microscope (IX83, Olympus, Singapore) using 543 nm continuous wave laser and 488 pulsed lasers for exciting hairpin and doubly labeled viruses, respectively. The beam was focused into the sample by a water immersion objective (× 60, NA 1.2; Olympus, Tokyo, Japan) after being reflected by a dichroic mirror (DM405/485/543/635 band pass, Olympus, Singapore) and the scanning unit. The fluorescence signal for hairpin and doubly labelled viruses were spectrally divided into donor and acceptor channels by a 650DCLP and 560 DCLP dichroic mirrors, respectively. The donor and acceptor fluorescence was recorded, by a set of single molecule avalanche photodiodes (SPADs) (SPCM-AQR-14, PerkinElmer Optoelectronics, Quebec, Canada), through a 615/45 and 670/40 band pass emission filter (Omega, VT), respectively for hairpin. On the other hand, virus’s fluorescence was collected through 513/17 and 615/45 band pass emission filter (Omega, VT), respectively. The intensities of the donor and acceptor channel are collated in 20 μs time bins in the SymPhoTime64 software and exported as text files. Using a MATLAB script, the proximity ratio is calculated for each time bin and the autocorrelation of the proximity ratio is calculated. The calculated autocorrelation of the proximity ratio is then fitted using the Levenberg-Marquardt iteration algorithm by Origin 9.1.

## Data availability

The datasets generated during and/or analyzed during the current study are available from the corresponding author on reasonable request

## Acknowledgements

The work and K.K.S. was supported by Singapore Ministry of Education Tier 3 grant (MOE2012-T3-1-008) awarded to T.W. and G.S.A.

## Author Contributions

K.K.S., and T.W. designed, analysed and interpreted the results. X.-X.L. performed the HDXMS experiments. K.K.S. and T.W. performed the TR-FRET and FRET-FCS experiments and wrote the section on TR-FRET and FRET-FCS. S.T. performed plaque assay and A.G. wrote software for FRET-FCS analysis. J.M. and P.B. provided computational data. X.Y.E.L. provided virus samples. K.K.S. and T.W. wrote manuscript and G.S.A., K.K.S., T.W. and P.B. contributed to manuscript revision.

